# Ecology and temporal dynamics of urban *Drosophila* species communities as potential indicators of biodiversity decline

**DOI:** 10.1101/2025.09.04.674250

**Authors:** Martin Kapun, Sonja Steindl, Maria Ricci, Manuel Löhnertz, Flora Strasser, Rui Qiang Chen, Lorin Timaeus, Nikola Szucsich, Elisabeth Haring

**Author notes:** co-correspondence. co-shared first authors (Martin Kapun, Sonja Steindl).

## Abstract

Understanding the impact of ecological factors on biodiversity is central in the context of accelerating climate change and biodiversity loss. Urban areas, as landscapes under particularly strong anthropogenic influence, are undergoing rapid ecological change, yet the consequences for urban biodiversity and ecosystem functioning remain poorly understood. In this study, we focused on fruit flies of the genus *Drosophila* – a diverse group of dipterans with variable ecological niches and degrees of synanthropy – to investigate species composition and community ecology in the metropolitan area of Vienna, Austria. With the help of numerous citizen scientists, we have collected approximately 18,000 specimens through dense spatio-temporal sampling both indoors and outdoors of human dwellings. A total of 13 *Drosophila* species were identified, with communities dominated by widespread cosmopolitan synanthropic species. Among these, *D. mercatorum* and *D. virilis* represent novel records for Austria. Comparisons to a previous study from more than 30 years ago revealed that the species richness in Vienna was more than 50% lower than before and showed that formerly common species were potentially replaced by neobiots. We further assessed ecological niches by intersecting species abundance data with high-dimensional, high-resolution earth observation datasets, which revealed distinct ecological preferences among species. In particular, the neozoan *D. mercatorum* emerged as a highly synanthropic species, tightly confined to urban areas with high levels of imperviousness. In summary, our study underpins the versatility of the *Drosophila* system as indicators of biodiversity loss in a rapidly changing world.

## Introduction

The genus *Drosophila* comprises over 1,600 described species of small dipterans, exhibiting an exceptionally wide range of ecological niches, behaviors and morphologies (Bächli, 1982; O’Grady and DeSalle, 2018). Their ecological strategies range from specialists, often restricted to narrow geographic areas and specific food sources, to opportunistic generalists with broad – sometimes global – distributions (Bächli and Burla, 1985; Bächli and Pite, 1982; Brake and Bächli, 2008). A subset of these generalists, characterized by a close association with humans (“synanthropism”), are collectively referred to as the “cosmopolitan guild” (Atkinson and Shorrocks, 1977; Miller et al., 2017; Nunney, 1996). These species thrive in highly disturbed environments influenced by human activities and are among the most successful biological colonizers. Accordingly, many *Drosophila* species exhibit a remarkable expansion potential. The most prominent example is *Drosophila melanogaster*, one of the best-studied model organisms in genetics and developmental biology (Bilder and Irvine, 2017; Hales et al., 2015; Jennings, 2011). Originally, *D. melanogaster was* native to sub-Saharan Africa (Lachaise et al., 1988). Its ability to adapt to diverse climates, coupled with the reliance on fermenting fruit (Mansourian et al., 2018) and other decaying organic matter, facilitated the early association of this species with human settlements (Haudry et al., 2020; Keller, 2007; Markow, 2015). The spread of *D. melanogaster* likely began thousands of years ago alongside the rise of agriculture and the establishment of trade routes (Chen et al., 2024; Kapopoulou et al., 2020). By the late 19th and early 20th centuries, with the rapid expansion of global trade and transport, the species had colonized temperate and tropical regions worldwide (Arguello et al., 2019). Today it occurs on every continent except Antarctica. Several other species have similarly undergone anthropogenic expansion. For example, *Drosophila simulans*, the sister species of *D. melanogaster* (Capy and Gibert, 2004), *Drosophila suzukii*, an invasive agricultural pest (Adrion et al., 2014), and *Drosophila virilis* (Mirol et al., 2008) have achieved near-global distributions within just a few centuries or even decades, largely driven by human mobility. Because of their varying ecological specializations and responses to environmental disturbance, *Drosophila* species can serve as reliable bioindicators for climate change, habitat alteration, and anthropogenic impact (Parsons, 1991; Poppe et al., 2013). In particular, community composition often reflects disturbance intensity: With increasing environmental degradation or anthropogenic impact, opportunistic and commensal species become more dominant (Avondet et al., 2003; Shorrocks, 1977).

Urban landscapes are extreme examples of anthropogenically altered ecosystems (Vitousek et al., 2008). They are characterized by high habitat heterogeneity, fragmentation, extensive impervious surfaces, air pollution, and pronounced urban heat islands (Marzluff, 2008; Szulkin et al., 2020). Green areas as well as nutritional resources are patchy, human-generated, and often ephemeral, such as organic waste or fermenting fruit in markets, compost bins, or household kitchens (e.g., Hong et al., 2024). These conditions may offer unique opportunities for certain *Drosophila* species while excluding others. Urban areas might thus act both as refuges for commensal species and as barriers for specialists with narrow ecological tolerances restricted to natural and undisturbed environments. Yet, despite the long-recognized existence of synanthropic drosophilids, systematic studies of their occurrence in urban environments remain scarce (Avondet et al., 2003; Bombin and Reed, 2016; Garcia et al., 2012; Ramniwas et al., 2024; Ulmer et al., 2024). Key ecological questions remain unanswered: Which urban microhabitats do different species exploit? Are urban assemblages dominated exclusively by generalists and hemerophilous species, which prefer or thrive in habitats influenced by humans, or can specialists persist in certain niches? Do species compositions differ among areas subjected to varying degrees of urban pressure, such as city centers versus peri-urban green spaces? Moreover, species inventories for large cities are rarely updated; for example, the most recent survey of *Drosophila* in Vienna dates back to 1994 (Gross and Christian, 1994). Recent urban *Drosophila* surveys have been conducted in France (Ulmer et al., 2024), southern Brazil (Hochmüller et al., 2010) and in Moscow (Gornostaev et al., 2024, 2023), but most sampling sites in these studies were relatively natural habitats embedded in urban matrices, rather than heavily built-up areas. Consequently, we still lack a clear picture of *Drosophila* ecology within the most anthropogenically altered urban microhabitats – such as inside and around buildings, near waste disposal points, and in transport hubs – where commensal species may reach their highest densities. Furthermore, most previous studies have sampled flies along predefined transects rather than through quantitative random sampling across entire urban areas, potentially limiting the ability to detect general patterns in urban environments.

Even for common species, detailed ecological and biological data, as well as phylogenetic relationships within the genus (Robe et al., 2005), often remain incomplete. While broad geographic occurrence data are available for many species, fine-scale distributional records – particularly in urban areas – are rare. In Austria, for example, GBIF lists 17 *Drosophila* species (DOI:10.15468/dl.22p76q), representing about 60% of the 27 species reported for the country. Of the 157 GBIF records, the vast majority originate from the citizen science platform iNaturalist, with additional contributions from INSDC Sequences, iBOL/BOLD, the Biodiversitätsdatenbank Nationalpark Donau-Auen, Observation.org, and the Biodiversitätsdatenbank Salzburg. Similar species inventories are found across Central Europe: Switzerland – 35 species (Merz and Schweizer Zentrum für die Kartographie der Fauna, 1998), Austria – 27 (Szucsich, unpubl.; Franz, 1989), Germany – 30 (AK-Diptera and Bächli, 2023), Czech Republic – 29 (Máca, 2009; Máca et al., 2015), Slovakia – 26 (Máca, 2009; https://gd.eppo.int/reporting/article-3303), Denmark – 23, Sweden – 26, Norway – 19, Finland – 27 (Bächli et al., 2004). The present study investigates *Drosophila* species distributions in urban habitats, with particular emphasis on occurrences inside and around buildings. Conducted within the framework of the citizen science project “Vienna City Fly”, this work aims to document the urban *Drosophila* species spectrum and to assess their ecological niches. This study is the first to focus predominantly on sampling flies within buildings, employing easily deployable funnel traps. In conjunction with outdoor collections, this approach enables a direct comparison of *Drosophila* communities between indoor and outdoor microhabitats. Specifically, we address the following questions: (1) Which *Drosophila* species occur in highly urbanized and rural areas in and around the city of Vienna, and what proportion are generalist versus specialist *Drosophila* species? (2) How does species composition vary with the degree of urbanization and microhabitat type? (3) Which environmental factors exert the greatest influence on species composition and abundance?

## Material & Methods

### Collection of flies

Collections of fly samples were carried out between July and December 2024 as part of a large-scale citizen science campaign entitled “Vienna City Fly” (https://nhmvienna.github.io/ViennaCityFly/), which involved 89 citizen science volunteers from the city of Vienna, but also from other Austrian federal states, who caught drosophilid vinegar flies using commercial traps that were placed within kitchens and/or gardens for a maximum of 14 days. The traps consisted of two parts, a translucent plastic container on the outside and a yellow plastic funnel with fine slits at the side to maintain airflow and three holes at the bottom of the funnel which allowed flies to enter the traps. Slices of fruit (e.g., banana or apple), which were placed at the bottom of the plastic container below the funnel, acted as bait and attracted flies into the trap. Behavioral constraints such as negative geotaxis prevented flies from escaping the trap. Caught specimens usually tried to leave the trap at the top of the plastic container which was sealed by the funnel and did not find the hole at the bottom of the funnel through which they initially had entered the trap. However, we observed that tiny individuals of species with overall small body size, such as *Drosophila melanogaster*, occasionally managed to escape through the air vents.

Throughout the sampling season, collectors returned or exchanged filled traps against new ones at the Natural History Museum of Vienna and provided information on collector’s ID (referring to the collecting site) and sampling duration. We explicitly requested additional information about the absence of flies, i.e. empty traps, and obtained this absence data from several collectors. The filled traps were then placed at –4°C for at least 30 minutes to anaesthetise the flies, prior to transferring all insects within a trap to Eppendorf tubes filled with 96% Ethanol. All collected flies were incorporated into the Diptera collection of the NHMW.

### Sampling area – the city of Vienna and its vicinity

We concentrated our sampling efforts in Vienna, a metropolitan area encompassing approximately 415 km² and comprising over 176,000 buildings (Bauer et al., 2024). The urban landscape included in our sampling is predominantly characterized by multi-family apartment buildings, including numerous public housing complexes, while single-family homes are more commonly found in peripheral districts. Although more than half of the total area of Vienna is classified as green space – including parks, forests, vineyards, and urban gardens – these are largely concentrated in the outer districts. The study primarily targeted the urbanized zones of the city, which present a highly heterogeneous mosaic of environments, ranging from densely built-up and impervious areas to green spaces and low-density residential neighborhoods. We also included samples collected in suburban or rural areas outside of Vienna, in the neighboring regions of Lower Austria and Burgenland, which are predominantly farmland and characterized by a humid continental climate (Köppen classification Dfb and Cfb; Köppen, 2011).

### Species identification

All sampled drosophilid flies were identified to species level under a light microscope following the identification key of Bächli & Burla (1985). We furthermore used additional literature to specifically distinguish species of the *repleta* group (Beuk, 1993). All identified specimens were then sorted, counted and stored in 96% Ethanol at 4°C for each trap and each species separately.

### DNA barcoding

For each *Drosophila* species, we further characterised several specimens from various localities by DNA barcoding using the mitochondrial (mt) cytochrome c oxidase subunit 1 gene (COX1) standard marker (Folmer et al., 1994). DNA extraction was performed in the DNA clean room of the Natural History Museum Vienna under strict contamination control protocols. The DNeasy Blood & Tissue Kit (Qiagen) was used for DNA extraction, following the manufacturer’s instructions, with a final elution volume of 50 µl. A negative control extraction was carried out without a DNA sample to detect potential contamination in the extraction reagents. The control extractions were later on included in the PCR reactions. All post-extraction work (thermocycling and post-PCR processing) were performed in a separate laboratory.

A partial region of the mitochondrial cytochrome c oxidase subunit 1 gene (COX1) was amplified using the PCR primers LCO1490 5’-GGTCAACAAATCATAAAGATATTGG-3’ and HCO2198 5’-TAAACTTCAGGGTGACCAAAAAATCA-3’ (Folmer et al. 1994) resulting in an amplicon 709 bp in length that was used as a genetic marker sequence (alignment length 658 bp). PCR reactions were conducted using the Multiplex PCR Kit (Qiagen, Hilden, Germany) in 25 μl reaction volumes containing 12.5 μl of Multiplex PCR Master Mix, 0.5 μM of each primer, and 2 μl of template DNA. The thermocycling conditions were as follows for all reactions: an initial denaturation at 94 °C for 15 minutes, followed by 40 cycles of 94 °C for 30 seconds, 52°C for 30 seconds, and 72 °C for 30 seconds, with a final extension at 72 °C for 10 minutes. Negative controls were included in all PCRs: one PCR reaction using the control extraction as a template and a PCR reaction with template DNA. PCR products were purified using the QIAGEN PCR Purification Kit and sequenced by SANGER sequencing in both directions using the original PCR primers (at Microsynth Austria).

The DNA barcodes generated contribute to the Austrian Barcode of Life (ABOL) initiative dedicated to record Austrian biodiversity. DNA barcodes can be found at the Barcode of Life database BOLD (https://v5.boldsystems.org) under the accession numbers XXX.

### Environmental variables

To identify potential links between *Drosophila* community composition and environmental factors, we obtained gridded climatic data from the data hub of GeoSphere Austria (https://data.hub.geosphere.at/) and demographic and administrative data from the data repository provided by the city of Vienna (https://www.data.gv.at/). Climatic data were obtained from two different sources: The SPARTACUS v. 2.0 dataset (Hiebl and Frei, 2016) from GeoSphere Austria consists of yearly (average temperature [TA; °C], total rainfall [RR; mm/m^2^] and total sunshine hours [SA; hours]), monthly (average temperature [TA; °C], total rainfall [RR; mm/m^2^] and total sunshine hours [SA; hours]) and daily (minimum daily temperature [TM; °C], maximum daily temperature [TX; °C], total precipitation [RR; mm/m^2^] and total sunshine hours [SA; hours]) datasets. The INCA dataset (Haiden et al., 2011) consists of hourly estimates of rainfall rate [RR; kg/m^2^], global radiation [GL; W/m^2^], relative humidity [RH2M; %], wind speed in eastward direction [UU; m/s], wind speed in northward direction [VV; m/s], dew point temperature [TD2M; °C] air temperature [T2M; °C] and air pressure [P0; Pa]. Both the INCA and the SPARTACUS datasets have a spatial resolution of 1km x 1km grid cells. We programmatically obtained the raw datafiles in NetCDF format from the datahub using R. Subsequently, we projected all layers to EPSG:4326 (WGS84) using the project() function of the R package terra (REF) and restricted all datasets to a geographical bounding-box (Longitude: 15.5°E – 16.7°E; Latitude: 47.5°N – 48.5°N) that includes the area covered by the city of Vienna and neighboring villages using the crop() function of the raster package in R. Using the resample function of the same package we further interpolated the gridded datasets to obtain a higher resolution raster of 100m x 100m grid cells for each continuous variable. For the yearly datasets, we extracted the single yearly value of the grid-cell closest to the latitude/longitude coordinate pair for each of the samples using the extract() function from the R package raster (Hijmans and van Etten, 2012). For the monthly datasets, we further extracted the values for the corresponding sampling month of the closest grid cell as described before. Finally, for the hourly and daily datasets we averaged all values in a 14-days interval prior to each collection date in the closest grid cells, since we assume that the weather conditions up to two weeks before the sampling may strongly influence the species composition in a given sample. We obtained land cover and land use data from the Austrian data portal (https://www.data.gv.at/) and from Copernicus Land Monitoring Service (CLMS, https://land.copernicus.eu/en). The Austrian data portal provides the land use map of Vienna, with 32 land use classes, in vector format. We defined a grid with 100m x 100m cells and for all cells computed the share of each class separately. We then projected the 32 raster layers thus obtained in EPSG:4326. We processed in the same way the “Protected Areas” layer. Similarly, we used the Buildings inventory of the city of Vienna to compute the share of built-up area and built-up volume in the 100m x 100m cells.

The CLMS Imperviousness density and Tree-cover density layers are raster layers providing the sealing resp. tree cover density in the range from 0% to 100% at 10m resolution. They were resampled at 100m x 100m and projected to EPSG:4326. In total we incorporated 58 independent variables into our analyses, including the information if flies were collected inside or outside of buildings, latitude, longitude and collection date of samples.

### Biodiversity data analyses

Based on the count data of each species, we first assessed the species-specific total abundance and average abundance of each species per trap. Furthermore, we assessed temporal variation by plotting the total abundance across the whole sampling season in two-weeks intervals for each of the identified species. To assess sampling completeness in the dataset, we performed species accumulation curve analysis using the vegan package in R (Dixon, 2003). Species accumulation curves were generated using random sampling methods, and asymptotic species richness was estimated by fitting Michaelis-Menten models to the accumulation data. Bootstrap confidence intervals (*n*=1,000 replicates) were calculated for total richness estimates, and sampling completeness was assessed using bootstrap Z-tests comparing observed richness to predicted asymptotic values. Statistical significance of sampling gaps was evaluated at *α*=0.05 to determine whether additional sampling effort would likely yield new species discoveries. Species data were aggregated by sampling locations and further separated by “indoors” or “outdoors” collections, with zero-abundance samples excluded. Subsequently, we calculated several estimators for *α*-diversity in each of the traps using the R package vegan. These estimators include the species richness (SR), i.e., the absolute number of species per trap, and the Shannon index (*H*’), which both estimate species richness and are thus strongly influenced by the number of rare species (Magurran, 2004). Furthermore, we calculated the Evenness (EV) and the Simpson diversity indices (1-*D*), which both summarize how evenly species are distributed in a sample (Magurran, 2004). We further estimated differences in community composition, i.e., *β*-diversity, among samples based on Bray-Curtis distances in R using the vegan package and employed non-metric multidimensional scaling (NMDS) to visualize the projected distances among the samples along the first two NMDS axes. We furthermore included species positions in NMDS plots, which are computed as weighted averages based on the abundances of species in the samples along the ordination coordinates. Subsequently, we scaled the 58 environmental variables with point estimates for each of the sampling sites in and around Vienna using the R function scale() and performed PCA using the PCA() function from the FactoMineR package (Lê et al., 2008). We then used the envfit() function from the vegan package in R to fit the first four PC-axes onto the NMDS ordination, using 99,999 permutations to assess the significance of correlations between the variables and the ordination axes.

### Ecological data analysis

For a more in-depth assessment of links between environmental variables and the abundance of *Drosophila* species, we employed Redundancy Analysis (RDA), which is a multivariate statistical method that combines elements of multiple regression and principal component analysis to explore and quantify the relationships between explanatory (i.e. environmental data) and response matrices (i.e., species abundance data). Since certain environmental datasets were not available for all sampling sites, we generated geographical subsets of the sampling data and analyzed these datasets separately. Before conducting the RDA, we tested for, and visualised intercorrelations among environmental factors in R using the cor() function and manually removed environmental factors including daily wind speed, global radiation, temperature, rainfall rate, monthly temperature means, and specific land use categories (outdoor sports, vineyard, water bodies, commercial areas) with high levels of positive or negative correlation (|*r*| > 90%) with at least one other factor and retained one of the intercorrelated factors. Subsequently, we standardized the values of the remaining environmental factors using the scale() function in R for z-score normalization and Hellinger-transformed the abundance data of the *Drosophila* species to reduce the impact of highly abundant species and excess zeros (e.g., Blanchet et al., 2014) while rendering the data appropriate for linear ordination methods such as principal component analysis (PCA) and redundancy analysis (RDA). We then used the rda() function from the vegan package in R to compute a full model including all environmental variables, as well as a null model conditioning on participant ID to control for collector effects. Subsequently, we performed forward model selection using ordiR2step() with 999,999 permutations to identify the combination of environmental variables that yielded the most informative model. To assess the significance of the most informative model and of individual terms, we conducted permutation tests with 9,999 iterations using the anova.cca() function from vegan. Finally, we visualized the RDA results using the ordiplot() function.

Furthermore, we performed species distribution modelling (SDM) using Random Forest algorithms to predict habitat suitability for *Drosophila* species across Vienna. We used the rast() function from the terra package (Hijmans et al., 2025) in R to combine land use, administrative boundaries, and annual average climate data (all in GeoTIFF format) into a comprehensive environmental predictor stack consisting of 44 distinct layers at a resolution of 230 x 300 grid cells, each of 100m^2^ size. We then aligned all raster layers to a common spatial template through resampling and cropped the whole stack to the study area extent. Species abundance data from the cleaned dataset restricted to Vienna were Hellinger-transformed to account for zero-inflation as described above, and spatial coordinates were converted to sf objects with an EPSG:4326 projection using the st_as_sf() function of the sf package (Pebesma and Bivand, 2023). Subsequently, we extracted environmental predictor values at each sampling location and trained Random Forest models for each of the 13 *Drosophila* species separately. We therefore employed the ranger package with 500 trees and mtry=3, using stratified sampling to partition the full dataset into 80% training and 20% testing subsets. Spatial coordinates and temporal variables were excluded from the predictor set to avoid potential data leakage and bias. Model performance was evaluated using *R*^2^, RMSE, and MAE on both training and test datasets for robust model assessment. We generated habitat suitability predictions across the entire study area and exported the results as GeoTIFF files. Results were visualized using ggplot2 with Stadia basemaps, displaying predicted abundance values with species-specific scaling limits (0 to 1 for the two most abundant species: *D. melanogaster* and *D. mercatorum*; 0 to 0.5 for the remaining species).

## Results

### Sampling and species abundance

During the sampling period, which spanned from late June to early December 2024, we received a total of 278 fly traps from 89 participants across the whole sampling area in Austria and 252 traps from 75 participants located in or near the city of Vienna, Austria (Figure 1). The majority of participants (*n*_all_=49 and *n*_Vienna_= 40) returned a single trap, and the remaining participants returned more, several of them even between 10 and 33 traps, throughout the sampling season resulting in an average of three traps per participant – both for the whole sampling area and for the city of Vienna, respectively. Samples outside of Vienna were collected in the Federal Provinces Burgenland (*n*=1) and Lower Austria (*n*=25).

**Figure 1.**
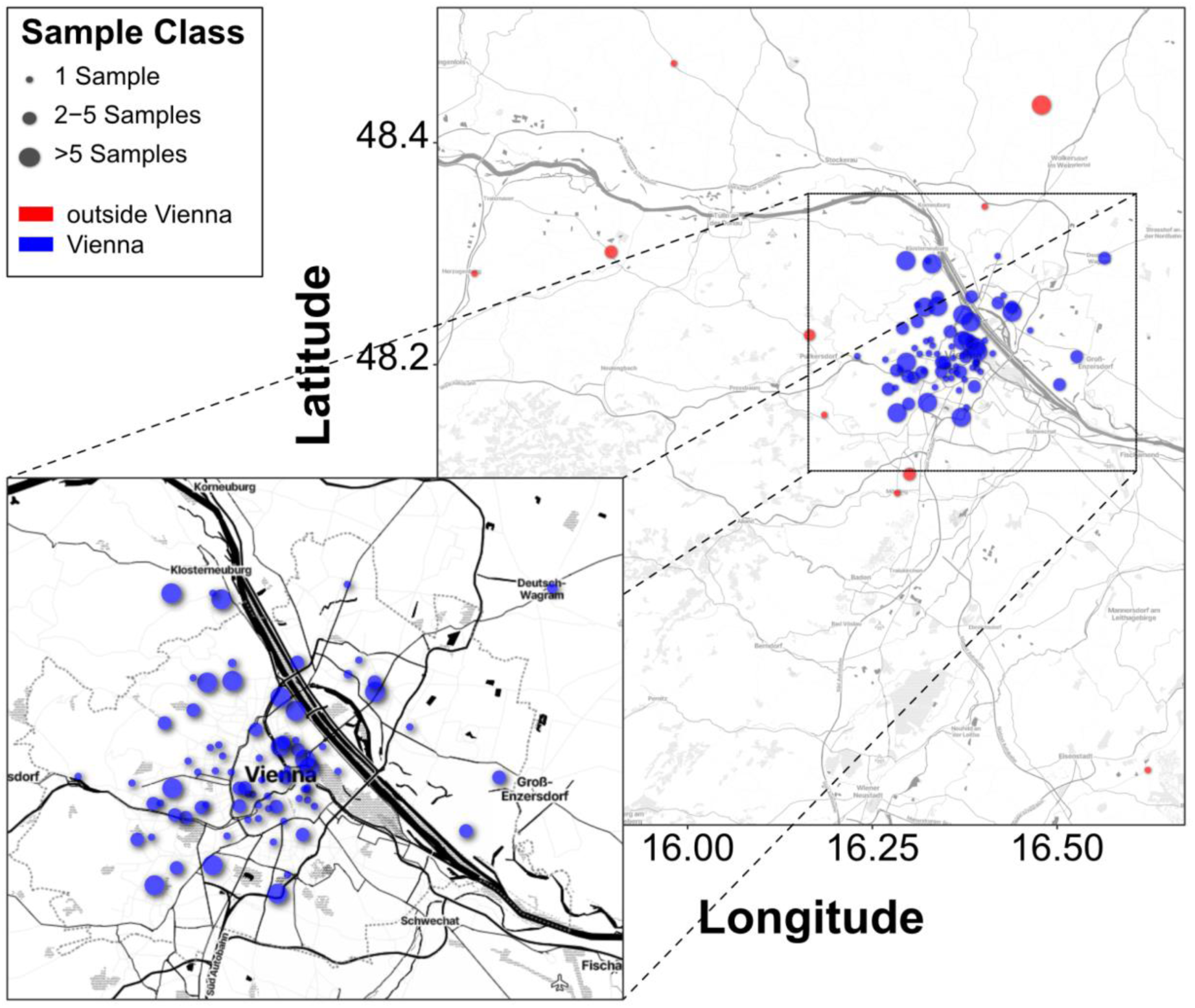
Sample distribution. Geographic distribution of sampling sites in Vienna and surrounding areas. Locations for which detailed land use and environmental data are available are highlighted in blue. Sampling sites outside this core area are shown in red. The size of the circle corresponds to the number of repeated sampling at a given location.

Our sampling effort yielded a total of 18,173 flies representing thirteen species of the genus *Drosophila* (Table 1 and Table S1). We identified several species of the *Sophophora* subgenus, including two members of the *melanogaster* group (*Drosophila melanogaster*, *Drosophila simulans*), one member of the *suzukii* group (*Drosophila suzukii*) and one member of the *obscura* group (*Drosophila subobscura)*. Furthermore, we found species of the subgenus *Drosophila* including *Drosophila virilis,* several species of the *immigrans*-*tripunctata* radiation lineage, including *Drosophila funebris*, *Drosophila immigrans*, *Drosophila phalerata*, and *Drosophila testacea* and members of the *repleta* species group including *Drosophila hydei*, *Drosophila mercatorum* and *Drosophila repleta* as well as one species from the subgenus *Dorsilopha* (*Drosophila busckii*).

**Table 1.** *Drosophila* species identified in this study. The final column indicates the number of samples from which DNA barcodes were obtained for each species.

| Subgenus | Species group | Species | Total Abundance | Sequenced |
| --- | --- | --- | --- | --- |
| *Drosophila* | *funnebris* | *D. funnebris* | 185 | 5 |
|  | *immigrans* | *D. immigrans* | 76 | 5 |
|  | *quinaria* | *D. phalerata* | 22 | 3 |
|  | *hydei* | *D. hydei* | 908 | 6 |
|  | *repleta* | *D. repleta* | 656 | 5 |
|  |  | *D. mercatorum* | 8851 | 6 |
|  | *testacea* | *D. testacea* | 44 | 3 |
|  | *virilis* | *D. virilis* | 90 | 3 |
| *Sophophora* | *melanogaster* | *D. melanogaster* | 6726 | 3 |
|  |  | *D. simulans* | 393 | 5 |
|  |  | *D. suzukii* | 195 | 3 |
|  | *obscura* | *D. subobscura* | 5 | 4 |
| *Dorsilopha* |  | *D. busckii* | 22 | 4 |

Eleven species were found in Vienna as well as in some surrounding localities. Only two species, *D. phalerata* and *D. virilis,* were found exclusively within the city borders. Species accumulation curve analysis indicated that sampling completeness was high (88.6%), with 13 observed species compared to a predicted total richness of 14.7 species (95% CI: 10.3-16.5). Bootstrapping indicated no statistically significant sampling gap (Bootstrap Z-test, *p*=0.336), which suggests that our funnel-trap sampling method captured most of the *Drosophila* species diversity present in the Vienna study area that is detectable with this approach.

To verify morphological species identifications, DNA barcoding was conducted on 2-6 individuals per species, in a total 55 individuals (Table 1). All sequences obtained, confirmed our morphological determinations. Along with corresponding reference specimens, these data contribute to the Austrian Barcode of Life initiative (ABOL; BOLD identification numbers: XXX).

To our knowledge, this study is the first to scientifically document the occurrence of two previously unrecorded species in Austria: *D. mercatorum* and *D. virilis*. One of these neozoans, *D. mercatorum* (*n*=8,851), as well as *D. melanogaster* (*n*=6,726) were the most frequent species (Figure 2A) in our collections. In contrast, *D. subobscura* was, by far, the rarest with only five individuals across all of the collections. In line with their overall abundance, *D. melanogaster* and *D. mercatorum* were consistently found in high numbers across many traps. Conversely, we found that certain species, such as *D. virilis* and *D. funebris* were overall rare but sampled in high numbers in only a few traps, as indicated by the large standard deviations shown in Figure 2B. We further observed that the location of the trap had a significant influence on the occurrence of certain species. While *D. suzukii*, *D. immigrans* and *D. simulans* were approximately 45, 43 and 11 times more likely to be caught outdoors, respectively, *D. mercatorum* and *D. repleta* were about 11 and 16 times more likely to be found indoors than outdoors (Table 2 and Figure 2C). The three species *D. phalerata*, *D. testacea* and *D. subobscura*, which were overall rare, were only caught outdoors

**Figure 2.**
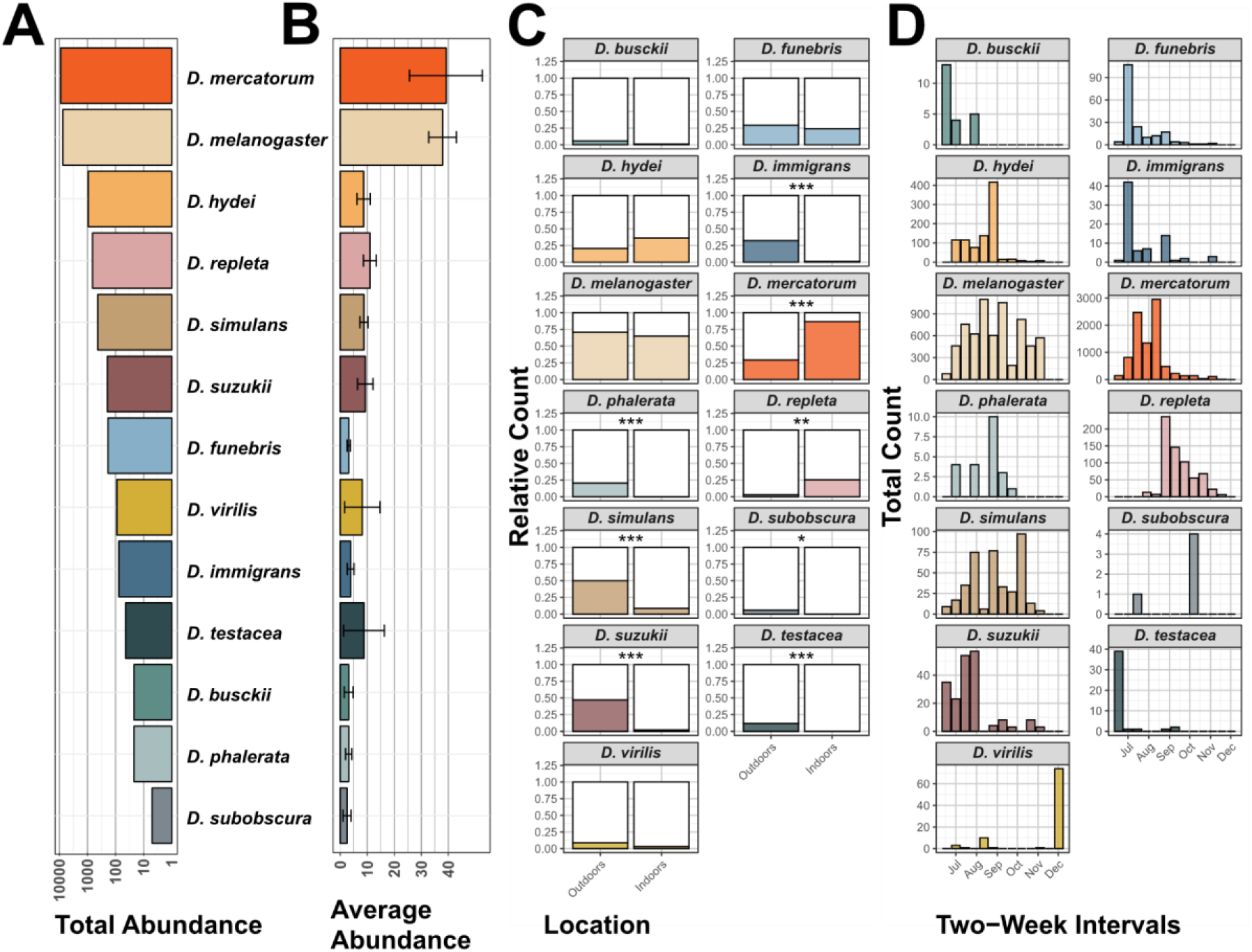
Abundance and phenology of the investigated *Drosophila* species. (A) Total number of individuals recorded per species across all sampling events. (B) Mean number of individuals per species per trap, with error bars representing standard deviations. (C) Proportion of traps (indoor vs. outdoor) in which each species was detected. Asterisks indicate significant differences in presence between trap locations (*=*p* < 0.05, **=*p* < 0.01, ***=*p* < 0.001). (D) Temporal distribution of species: histograms showing absolute counts per species in two-week intervals over the full sampling period.

**Table 2.**
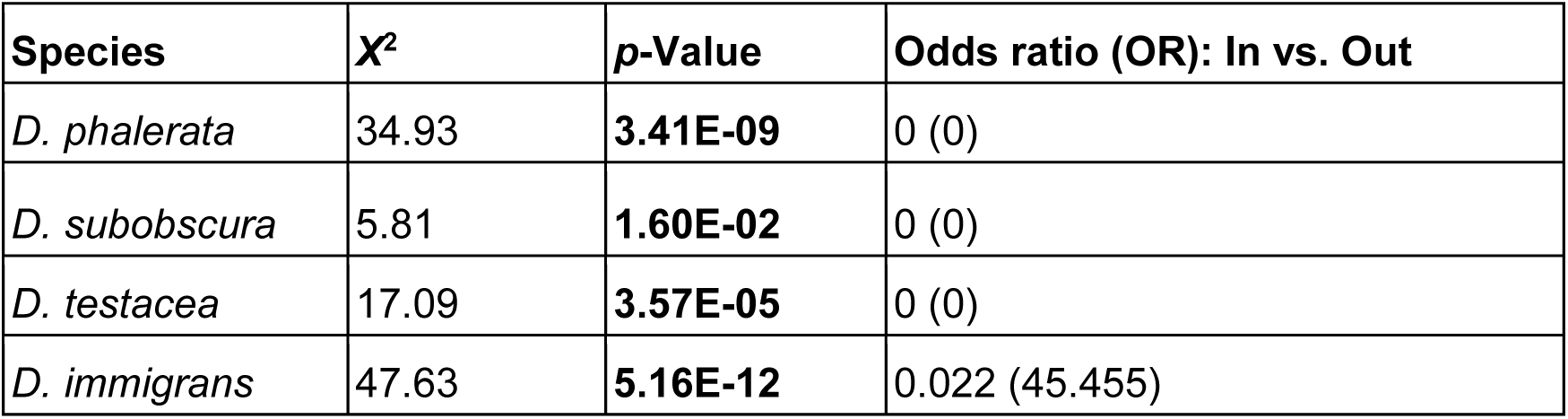
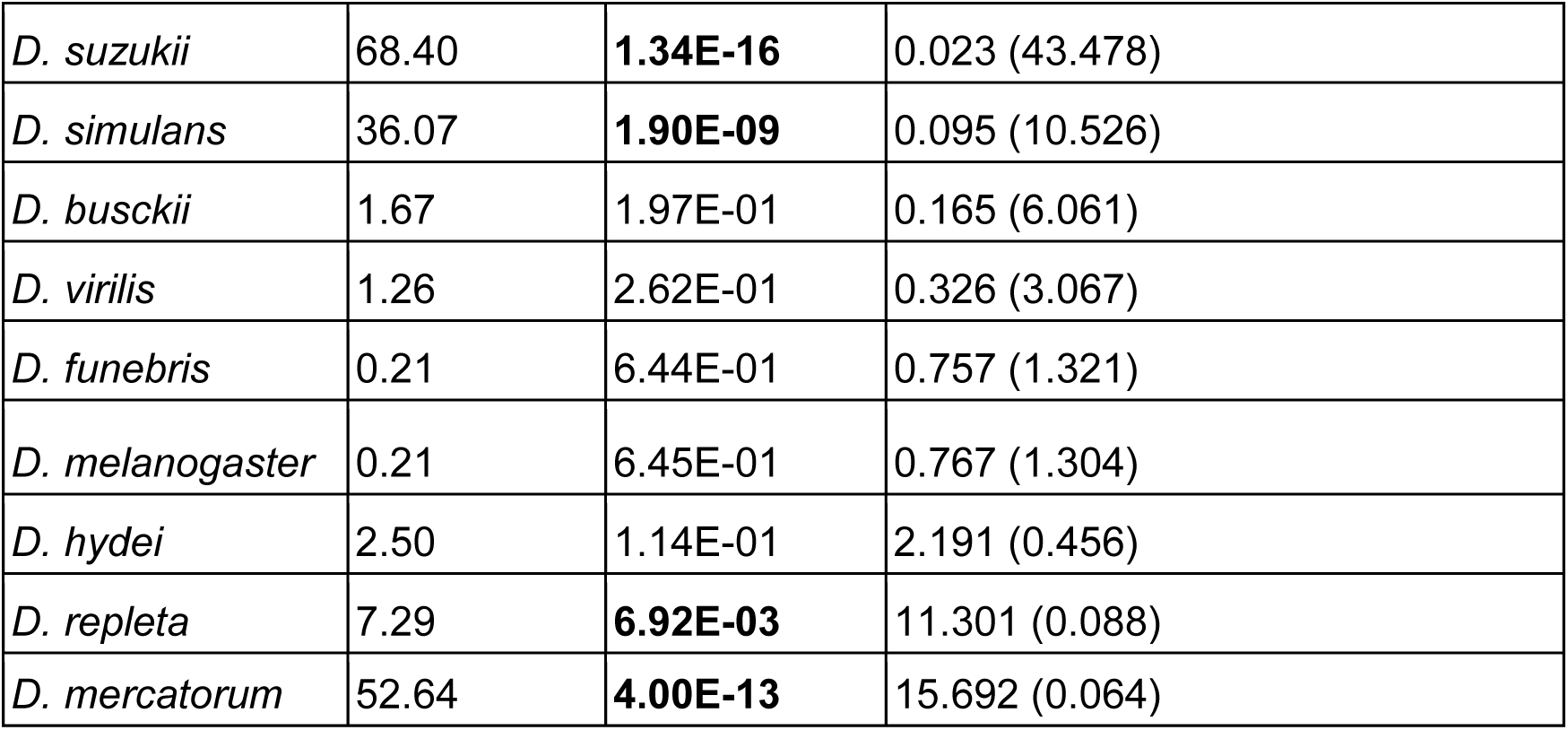
Results of Chi-square tests comparing the counts of samples collected indoors (*n*=233) and outdoors (*n*=45) for each species. The final column reports the odds ratio, reflecting the relative likelihood of a species being detected indoors compared to outdoors (or outdoors compared to indoors, as indicated in parentheses). Significant differences are highlighted in bold. Species with odd ratios of zero were only collected outdoors.

Furthermore, we observed differences in the phenology of the different species (Figure 2D). *D. busckii*, *D. funebris*, and *D. testacea* were predominantly trapped early in the year. Among the abundant species, *D. melanogaster* and *D. simulans* occurred commonly across the whole sampling season. Conversely, we found that *D. suzukii* was mostly caught early in the season, while *D. virilis* occurred only very late during the sampling season. However, we caution that our random sampling scheme at different locations and time-points may contribute to sampling bias that could influence the seasonality patterns observed here.

### Biodiversity and ecology

We next assessed *α*-diversity across samples and found that traps generally exhibited moderate levels of diversity (Figure 3A). The mean Shannon index (H′) was 0.59 (SD=±0.40), and the mean Simpson diversity index (1 − D) was 0.34 (SD=±0.23), reflecting moderate species diversity across the dataset. Species richness ranged from 1 to 9 species per trap, with a mean of 2.81 (SD=±1.36). The distribution of richness was positively skewed, indicating that most traps were dominated by just a few species, while only a minority harbored more diverse communities. Evenness was overall similarly moderate, with a mean of 0.66 (SD=±0.22), suggesting that, on average, no single taxon dominated the assemblages. However, unlike richness, evenness values were negatively skewed. This pattern indicates that most samples had relatively balanced species abundances, while a smaller subset of traps exhibited low evenness – likely due to one or a few species dominating the community, despite higher overall richness.

**Figure 3.**
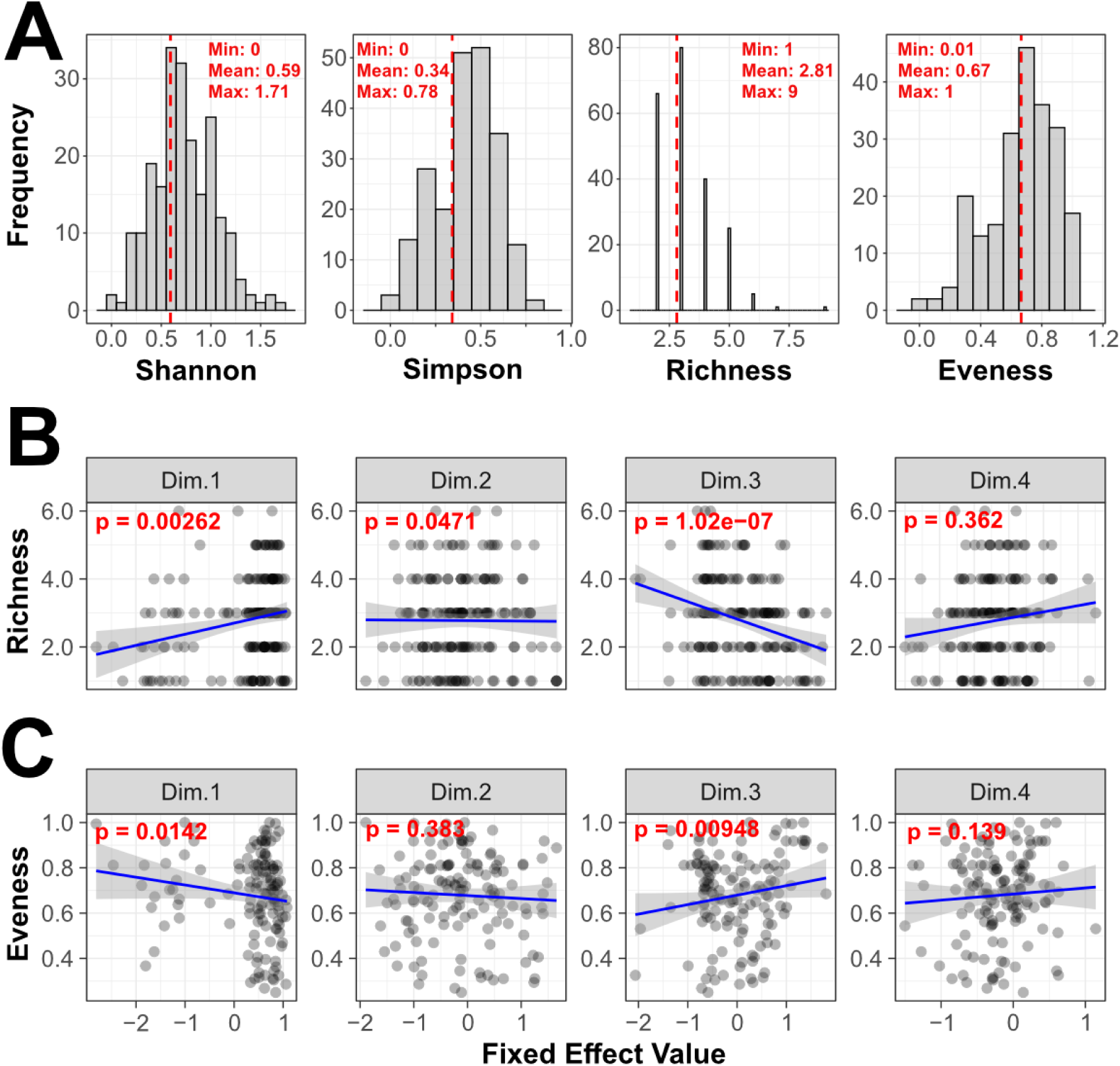
*α*-diversity and the influence of the environment on species diversity. (A) Histograms illustrating the distribution of α-diversity metrics across all traps. Insets display the minimum, mean, and maximum values, with vertical red dashed lines indicating the mean for each metric. The other two subpanels show correlations between Species Richness (B) and Evenness (C) and the first four principal components (PCs) from a PCA based on 58 environmental variables (see also Figure S1). P-values were derived from multiple ANOVA models, with α-diversity metrics as dependent variables and latitude, longitude, and the four PC axes as independent variables. The four PC-axes were dominated by short-term weather conditions, long-term climatic conditions, urbanization and aerial factors, such as wind and rainfall, respectively (see Table S3).

To test if environmental factors had a significant influence on *α*-diversity, we focused our analysis on samples collected within and around the city of Vienna, where detailed information on land use, climate and weather conditions at the sampling time-points were available. PCA of 58 environmental factors (Table S2), which were included in the analyses, revealed that the information of the first four axes (Dimensions) were dominated by (1) short-term weather conditions, (2) long-term climatic conditions, (3) anthropogenic influence, including land-use as well as (4) wind, precipitation and air pressure, respectively (Figure S1 and Table S3). Multiple linear regression revealed that Species Richness was significantly correlated with the first three principal component axes (Dim.1, Dim.2, and Dim.3; Figure 3B; Table S4). Notably, we found a strong negative correlation with Dim.3, which is primarily associated with anthropogenic factors. This suggests that Species Richness declines as urbanization and human influence increase. Moreover, the positive correlation with Dim.1 indicates that Species Richness increases with lower average annual temperature, higher average precipitation and altitude. We likewise found significant correlations of Dim.3 with the Shannon and the Simpson diversity indices (Table S4). Conversely, we found the opposite correlation patterns for Eveness, suggesting that urban areas are dominated by a few, evenly distributed *Drosophila* species (Figure 3C; Table S4).

We further assessed *β*-diversity by calculating pairwise Bray-Curtis distances among all samples. The subsequent ordination with NDMS and correlation of species and PC axes from the environmental data with the two NMDS axes revealed substantial species-specific differences in the occurrence among the traps. Along the first NMDS axis we observed a strong gradient ranging from *D. mercatorum* and *D. repleta*, which we previously identified to predominantly occur indoors (see above), to *D. suzukii*, *D. phalerata*, and *D. testacea*, which all showed a strong preference for outdoor habitats in our dataset (Figure 4A). Consistent with this observation, we found that Dim.3 of the environmental PCA, which was influenced by environmental factors associated with land use and human activity, was most significantly associated with the ordination space, Dim.3 explained about 14.3% of the variation in the ordination, which was statistically highly significant (*p* < 1e-5; Table S5). The remaining three PC axes each explained 1.6% (Dim.1; *p* > 0.05), 2.11% (Dim.4; *p* > 0.05) and 2.79% (Dim.2; *p=*0.044) and thus had either no, or only very small influence on the species diversity. While not the dominant drivers of the variation, this result further indicates a substantial influence of urbanization on community composition and species distribution.

**Figure 4.**
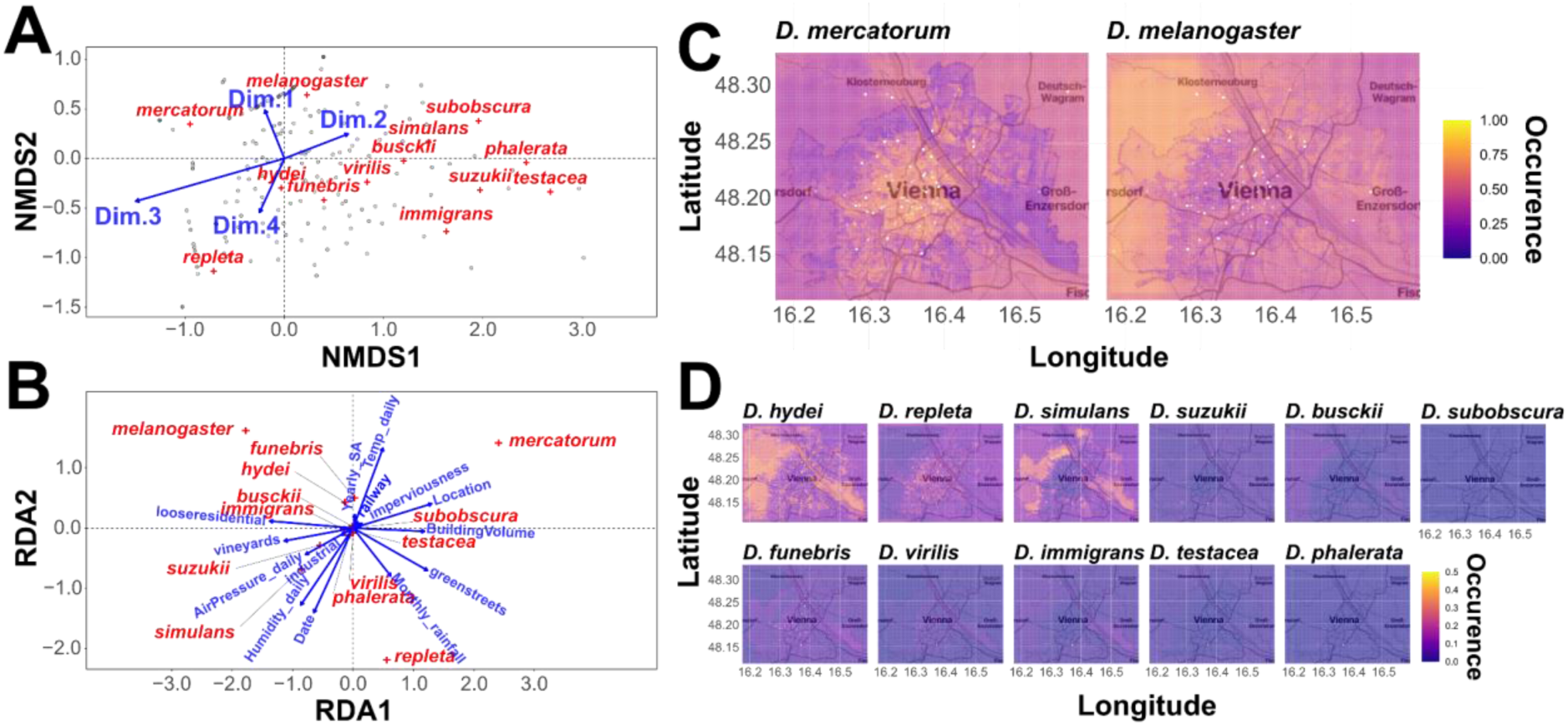
*β*-diversity and the influence of environmental factors on species distribution. (A) NMDS plot based on Bray-Curtis dissimilarities, with grey dots representing individual traps. Red crosses indicate the average position of each *Drosophila* species along the two NMDS axes, while blue arrows represent correlations with the first four principal components (PCs) from a PCA of 58 environmental variables (see also Figure S1). (B) Redundancy Analysis (RDA) plot showing the first two canonical axes. Red crosses indicate species’ positions in RDA space, and blue arrows represent correlations with 14 environmental variables selected by forward regression. (C) and (D) display results from species distribution models based on random forest modelling. Color gradients indicate predicted occurrence probabilities, ranging from 0 to 1 in panel C, and from 0 to 0.5 in panel D (see also Table 3).

**Table 3.** Performance metrics of species distribution models based on a random forest approach used to predict species-level abundance across training and test datasets. Reported metrics include R² (coefficient of determination) and RMSE (Root Mean Square Error) for both training and test sets, as well as MAE (Mean Absolute Error) for the test set. While the Root Mean Square Error (RMSE) measures the average size of prediction errors and gives more weight to larger errors by squaring them before averaging and then taking the square root, the Mean Absolute Error (MAE) quantifies the average absolute difference between predicted and observed species abundance, indicating overall prediction accuracy regardless of whether the model over-or under-predicts the abundance. NA values indicate species for which no test data were available due to limited sample sizes.

| Species | Train $R^2$ | Train RMSE | Test $R^2$ | Test RMSE | MAE |
| --- | --- | --- | --- | --- | --- |
| *D. mercatorum* | 0.567 | 0.238 | 0.461 | 0.270 | 0.202 |
| *D. melanogaster* | 0.396 | 0.298 | 0.210 | 0.334 | 0.293 |
| *D. virilis* | 0.591 | 0.055 | 0.133 | 0.040 | 0.009 |
| *D. immigrans* | 0.182 | 0.068 | 0.118 | 0.067 | 0.024 |
| *D. hydei* | 0.498 | 0.125 | 0.104 | 0.183 | 0.128 |
| *D. simulans* | 0.399 | 0.164 | 0.062 | 0.175 | 0.100 |
| *D. funebris* | 0.321 | 0.094 | 0.057 | 0.121 | 0.075 |
| *D. repleta* | 0.504 | 0.190 | 0.022 | 0.267 | 0.185 |
| *D. suzukii* | 0.295 | 0.139 | 0.004 | 0.091 | 0.041 |
| *D. busckii* | 0.554 | 0.016 | 0.002 | 0.023 | 0.004 |
| *D. subobscura* | 0.544 | 0.013 | NA | 0.006 | 0.002 |
| *D. phalerata* | 0.098 | 0.066 | NA | 0.018 | 0.007 |
| *D. testacea* | 0.081 | 0.027 | NA | 0.008 | 0.003 |

Complementing the previous analyses, we applied redundancy analysis (RDA) to assess the influence of specific environmental factors on species composition. Using forward selection, we identified 14 environmental variables that contributed to the best-fitting model, which was constrained by latitude and longitude (Table S6). This model explained approximately 26.9% of the variation in species composition (adjusted *R*²=0.2690) and the most significant factors were “Location” (“indoors/outdoors”; *F*_1,213_=21.74, *p* < 0.001), “sampling date” (*F*_1,213_=20.3, *p*<0.001), the proportion of greened road space (“greenstreets”; *F*_1,213_=11.98, *p*<0.001), the building volume in a grid cell (“BuildingVolume”; *F*_1,213_=9.69, *p*<0.001), the proportion of loosely built residential areas (“looseresidential”; *F*_1,213_=7.84, *p*<0.001) and monthly precipitation (“Monthly_RR”; *F*_1,213=_5.66, *p=*0.002). In the RDA plot (Figure 4B), species positioned at the periphery typically occurred across many samples, while those near the center were often rare and observed in only a few samples. Notably, the two most common species, *D. melanogaster* and *D. mercatorum*, were located at opposite ends of the first RDA axis but shared similar positions along the second axis. In contrast, *D. repleta* appeared at the opposite end of the second axis, suggesting ecological differences among these three species. Most of the other species clustered near the center of the plot, reflecting their broader and more uniform distribution across environmental gradients or their rareness.

Consistent with the results above, we found that the distribution of *D. mercatorum* on the RDA plot was positively correlated with several environmental factors, such as sampling location, building volume, and daily temperature. This is indicated by arrows pointing toward the species’ position on the RDA plot along the first or second RDA axis, providing further evidence that *D. mercatorum* has a strong ecological preference for habitats with a high anthropogenic influence and elevated daily temperature. The second most common species, *D. melanogaster*, which is commonly assumed to be an obligatory human commensal, appears to be likewise, but less strongly influenced by temperature. However, in contrast to *D. mercatorum* this species seems to be negatively correlated with building volume but more common in loosely built residential areas, which may suggest that it prefers less urban but rather sub– and peri-urban habitats. Both *D. simulans* and *D. repleta*, which represent less common, but still abundant species, were positively influenced by monthly rainfall and daily humidity but negatively correlated with daily temperature, which may indicate that these species prefer a cooler and more humid climate than the two aforementioned species. However, we caution that these analyses may be confounded by the unequal number of samples provided by citizen scientists, which may bias towards locations that have been sampled multiple times. To test the robustness of our results, we repeated the RDA analysis after removing daily and monthly climatic variables and collapsing all samples collected at the same sampling site. The collapsed abundance data were Hellinger-transformed to account for unequal sample sizes. Consistent with the previous results, we found a qualitatively similar distribution of *D. mercatorum*, *D. melanogaster*, and *D. simulans* in the RDA plot (Figure S2). The main differences were that *D. repleta* was located closer to *D. mercatorum*, and *D. hydei* was positioned on the opposite side of the second RDA axis, relative to *D. mercatorum*. Moreover, *D. repleta*, and *D. melanogaster* were farther from the plot center. Furthermore, we again found that “Location” (indoors/outdoors; *F*_1,74_=7.04, *p* < 0.001) and building volume per grid cell (“BuildingVolume”; *F*_1,74_=4.26, *p*=0.01) were significantly associated with the distribution of *D. mercatorum* (Table S7). Overall, the model explained 14.6% of the variance in species distribution (adjusted *R*²=0.14581). We therefore consider our main findings from the full model to be robust.

We finally performed species distribution modelling (SDM) based on random forest models using the abundance dataset collected from within and around Vienna and integrated gridded datasets available for 48 environmental factors (Table S2). Due to their high overall abundance, we specifically focused on the two species, *D. melanogaster* and *D. mercatorum*. The species distribution model performed particularly well for *D. mercatorum*, with a high test *R*² of 0.461 and low error metrics (RMSE=0.27, MAE=0.202; Table 3), indicating strong predictive power and a good fit to the data. In contrast, the model for *D. melanogaster* showed rather poor performance, with a low test *R*² of 0.149 and higher error rates. These findings suggest that *D. mercatorum*’s distribution is more tightly linked to the measured environmental factors, while *D. melanogaster* may be influenced by additional variables not captured in the model. The SDM plot shown in the left panel of Figure 4C confirms the results from the previous analysis indicating that *D. mercatorum* is strongly confined to urban areas with high levels of imperviousness and anthropogenic influence and showing that *D. melanogaster* commonly occurs across a broad range of habitats within and around the city of Vienna. Among the other species in our dataset*, D. virilis* also showed reasonably good performance (test *R*²=0.133, RMSE=0.040). However, most other species, including *D. simulans*, *D. busckii*, *D. repleta*, and *D. suzukii*, exhibited very low or near-zero test *R*² values despite low error metrics in some cases (Figure 4D), which is potentially due to the low sample sizes in some of the investigated species. This pattern further suggests that the models for these species likely overfitted to the training data and did not generalize well.

## Discussion

### The persistence and influence of urbanity on *Drosophila* communities

Urban landscapes are often considered ephemeral environments with a high turnover rate that negatively affect biodiversity, while favoring ecological generalists and neobiota (Johnson and Munshi-South, 2017; Lewthwaite et al., 2024; McGlynn et al., 2019; Sukopp, 2008; Szulkin et al., 2020). To investigate these ecological dynamics and better understand the impact of human activity on biodiversity in urban landscapes, we analyzed the composition of *Drosophila* species communities in the metropolitan area of Vienna in Austria. Specifically, we focused on samples collected in close proximity to human settlements, both indoors (within kitchens, living or dining rooms) and outdoors (on balconies and in gardens).

Consistent with previous studies (for a review, see Sukopp, 2008) and our expectations, we found that *Drosophila* species richness was negatively correlated with urbanity. The *Drosophila* community of the Vienna city area was dominated by synanthropic and cosmopolitan members of the “cosmopolitan guild”, namely *D. melanogaster*, *D. simulans*, *D. funebris*, *D. hydei*, *D. immigrans* and *D. repleta* and a few specimens of *D. busckii*. As shown by previous studies, for example, in France (Ulmer et al., 2024), North America (Avondet et al., 2003; Bombin and Reed, 2016) and with the exception of *D. funebris* also in South America (Ferreira and Tidon, 2005; Garcia et al., 2012; Gottschalk et al., 2007), these species have been found to be common in world-wide urban areas. This highlights the strong synanthropic and hemerophilous character of these species and underscores their ability to thrive in highly disturbed environments, such as urban habitats, even across different climatic zones (Atkinson and Shorrocks, 1977; Nunney, 1996).

We also detected recent additions to the neobiotic *Drosophila* fauna of Vienna, such as *D. suzukii*, a fruit pest originating from Asia that was first reported in Austria in 2012 (EPPO Global Database [https://gd.eppo.int/]; Asplen et al., 2015). In addition, we found two species not previously reported in the scientific literature to occur in Austria to date: *D. mercatorum* and *D. virilis*. *Drosophila mercatorum*, which was the most common species in our collection, was originally described from California and is distributed throughout the United States, Mexico, and South America. It has been introduced to Europe in the last century, first recorded in Spain (Prevosti, 1953). Subsequent reports from various countries – from Portugal to Ukraine – document the continued expansion of its range (Adaschkiewitz and Gossner, 2013; Ivannikov and Zakharov, 2000; Kraaijeveld, 1992; Pité, 1972; Widmann and Bächli, 2022). In Austria, a few records on the citizen-science platform iNaturalist and a sample barcoded by L. Timaeus already in 2022 (ABOL-BioBlitz22-0415; https://portal.boldsystems.org/record/TDAOE2549-23) indicate that the species’ presence was not entirely unnoticed. However, as *D. mercatorum* is easily confused with *D. repleta*, it may have remained overlooked in many locations for a longer time. For instance, Vilela (1983) reported this species from Zimbabwe based on a specimen originally collected in 1935. *Drosophila virilis*, the second newly reported species for Austria which was rather rare in our collections, originated in Asia – likely from forested regions of China or the arid zones of Iran and Afghanistan (Throckmorton, 1982) – and only recently expanded its range as a human commensal, now occurring widely across the Northern Hemisphere (Mirol et al., 2008; Vieira and Charlesworth, 1999; Wang et al., 2006).

We failed to identify several *Drosophila* species that have been previously detected in the Vienna area 34 years ago based on a similar sampling design (Gross and Christian, 1994). Only 10 species – *D. busckii*, *D. funebris*, *D. hydei*, *D. immigrans*, *D. melanogaster*, *D. phalerata*, *D. repleta*, *D. simulans*, *D. subobscura*, and *D. testacea* – were also identified in our survey. In total, 12 species reported by Gross & Christian (1994) were not detected in the present study. Among them were nine *Drosophila* species – *D. ambigua*, *D. bifasciata*, *D. confusa*, *D. deflexa*, *D. helvetica*, *D. kuntzei*, *D. limbata*, *D. obscura*, *D. rufifrons*, *D. subsilvestris*, *D. tristis*, and *D. tsigana* – as well as three taxa that are no longer classified within the genus *Drosophila*. *D. rufifrons* and *D. deflexa* are now placed in *Scaptodrosophila* (Grimaldi, 1990), while *D. confusa* is assigned to *Hirtodrosophila* (Grimaldi, 1990). Thus, nine *Drosophila* species appear to be absent, which corresponds to a 47.4% *Drosophila* species reduction over time. Two of these belong to the *quinaria* group, and one to the *melanica* group. The most dramatic changes were observed in the *obscura* group. Seven members of the *obscura* group were not observed in the present study. These include *D. obscura*, a species native to Europe and reported as very common by Gross & Christian (1994).

Moreover, *D. subobscura*, which was the most common species in the previous study, was the rarest in our survey, represented by only five specimens. We further found that *D. deflexa* and *D. tsigana* (Burla and Gloor, 1952) – two *Drosophila* species that were previously detected for the first time in Austria or even in Central Europe (Gross and Christian, 1994) – were absent from our collections which may suggest that these species failed to establish in the study area.

In summary, the comparison of species composition over the past three decades reveals a dramatic turnover and reduction in *Drosophila* diversity in the metropolitan area of Vienna, consistent with the accelerating global biodiversity loss observed in recent decades (Drenckhahn et al., 2020; Isbell et al., 2023). The reasons for these changes are likely manifold: (1) Driven by global climate change, the average annual temperature in the inner city and the Vienna catchment area has risen by more than 2 °C during this period (see Figure 5), which may have reduced the competitiveness of *Drosophila* species with narrow temperature niches (Capek et al., 2025; Kim et al., 2020; Sillero et al., 2014). (2) Furthermore, increasing temperatures may have triggered indirect ecological effects, such as shifts in food availability or changes in the prevalence of parasites, parasitoids and insect diseases, which could differentially affect the *Drosophila* community (Corcos et al., 2019). (3) Alterations in human activities, such as waste management practices or the use of insecticides, may have disproportionately impacted the species that are now absent compared to those still detected in our survey (e.g., Collins et al., 2024). (4) Finally, competition with newly introduced neobiotic *Drosophila* species occupying overlapping niches may have contributed to the decline of formerly common species (Krijger et al., 2001; Rombaut et al., 2023). In addition to these biological and ecological explanations, we cannot exclude the possibility that differences in sampling design also contributed to the observed patterns. In contrast to Gross & Christian (1994), who predominantly collected their 28 spatio-temporal samples outdoors, the majority of our samples (83.8%, *n*=233) were collected indoors, with only 45 samples obtained outdoors. Moreover, Gross & Christian (1994) placed their traps in more rural and semi-natural landscapes, which may have facilitated the capture of less synanthropic species compared to our study. However, seasonal differences, bait choice and subtle differences in the trap design may have favoured or repelled different species in this study and the former fly collections. We therefore speculate that the observed differences in species abundance and community composition may reflect major ecological and biological changes, potentially reinforced by differences in sampling methods between the present and the previous study by Gross & Christian (1994).

**Figure 5.**
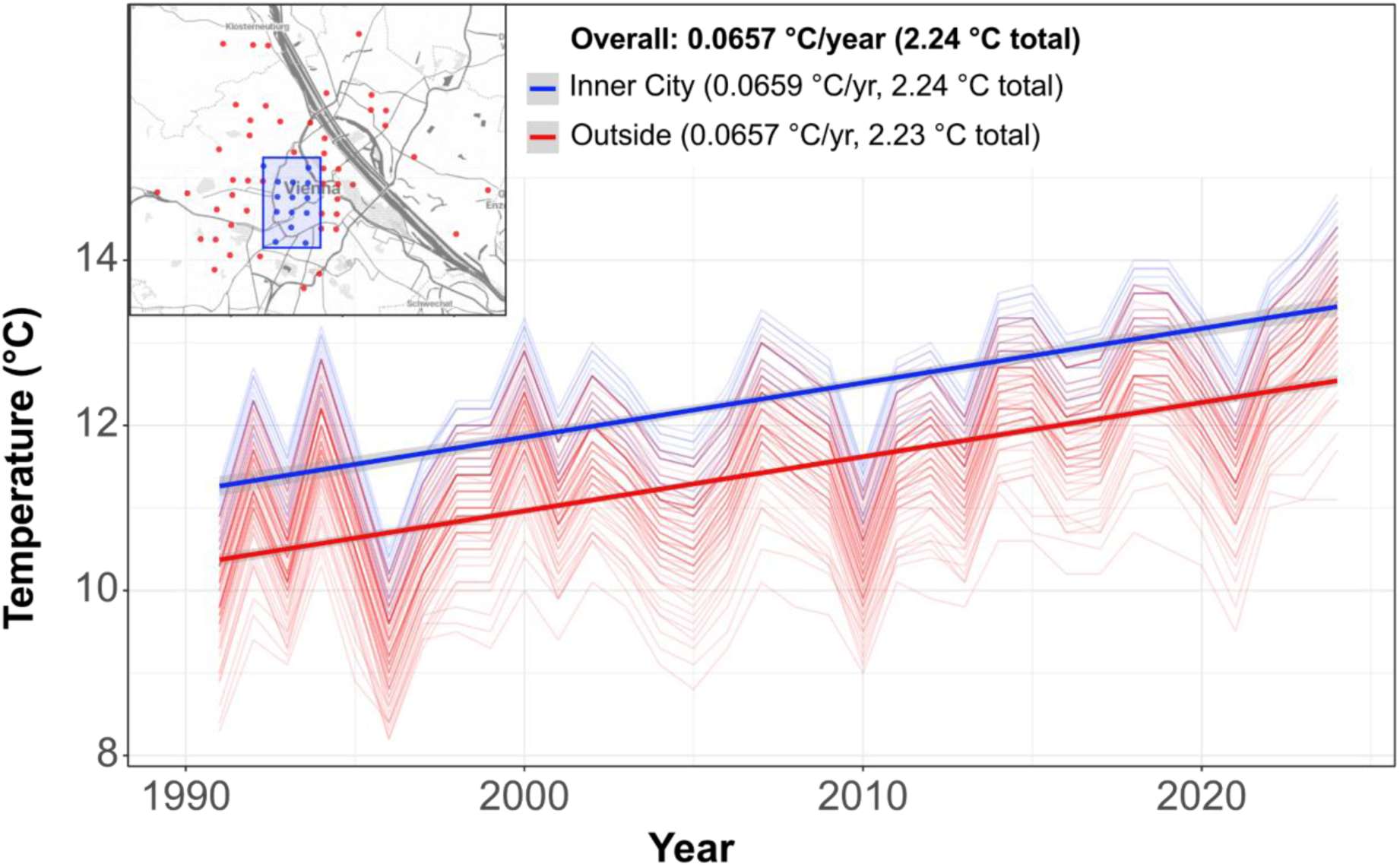
Annual average temperature change in Vienna. Line plot depicting annual average temperatures at the sampling locations since 1991. Blue lines represent samples from the central urban region with a high degree of imperviousness (highlighted by a blue box in the map at the top left). Red lines represent the remaining sampling locations. Bold straight lines indicate regression lines from linear models calculated separately for the two groups. Insets at the top show the average annual temperature increase as well as the total increase across the whole time span, both for the full dataset and for the two subsets.

### The ecology of urban and suburban *Drosophila*

Differences in ecological niches can strongly influence the composition of *Drosophila* communities across environments. Previous studies have shown that many *Drosophila* species, even within the cosmopolitan guild that commonly coexist sympatrically, exhibit substantial differences in their food preferences and breeding sites. For example, *D. melanogaster*, *D. simulans*, and *D. suzukii* – all members of the *Sophophora* subgenus-prefer fruits at different stages of decay as food sources (Atkinson and Shorrocks, 1977; Brake and Bächli, 2008). In contrast, species of the *obscura* group are typically associated with forest habitats and are thought to rely on fungal substrates, although detailed information on their breeding sites remains scarce (e.g., Kimura, 1980). Members of the *quinaria* group, such as *D. phalerata* and *D. testacea* (Scott Chialvo et al., 2019) are recognized mushroom feeders, whereas *D. virilis* preferentially feeds from slime flux of trees but also breed on other substrates (Spieth, 1979). Meanwhile, members of the *repleta* group, such as *D. repleta* and *D. mercatorum*, utilize decaying plant material, including rotting cacti in their native habitats in the Americas (Hasson et al., 1992; Oliveira et al., 2012).

Because the traps provided by citizen scientists contained a wide variety of baits – including citrus fruits, berries, apples, bananas, and other fruits – it was not possible to directly assess the food preferences of the investigated fruit fly species in this study. To account for the fact that ecological niches are shaped by more than food sources alone, we complemented our survey with a comprehensive environmental dataset comprising climatic, weather, and land-use variables at the sampling locations to analyze the ecological preferences of the observed *Drosophila* species.

Complementary analyses based on redundancy analysis and environmental correlations with Bray–Curtis distances revealed that short-term weather conditions, such as daily temperature and rainfall, strongly influenced species composition and abundance. However, the strongest impact was associated with the degree of urbanization, measured, for example, by land-use types as well as by building volume and area at the sampling sites.

The neozoan fly *D. mercatorum*, and to a lesser degree *D. repleta*, clearly stood out as isoanthropic species. In particular, *D. mercatorum* was rarely captured outdoors and was found predominantly in highly urbanized regions with the greatest levels of imperviousness. By contrast, *D. melanogaster* – commonly regarded as a strict human commensal with a high degree of hemerophily – showed a preference for more suburban areas with substantial green space. Our analyses further indicated marked ecological differences between *D. melanogaster* and its sister species *D. simulans*, the latter being found mainly in rural residential areas and only rarely indoors. At the opposite end of the spectrum, we observed species such as *D. phalerata* and *D. testacea*, which are considered native to Central Europe and were never detected indoors or in highly impervious areas.

In summary, our analyses provide evidence for highly distinct ecological niches among most of the investigated species. This is particularly remarkable given that the majority of specimens in this study were collected indoors and thus only indirectly exposed to weather and climatic conditions. Our findings therefore suggest that flies are highly mobile and primarily transient visitors to human dwellings, spending much of their life cycle outdoors where they are directly influenced by the environmental factors shaping their niches.

## Conclusion

In this study, we comprehensively investigated the *Drosophila* community through dense spatiotemporal sampling in and around the metropolitan area of Vienna, Austria – a region characterized by a diverse range of landscapes and land-use types. Our results highlight the power of citizen science for biodiversity assessment, made possible through the coordinated efforts of many contributors who collected samples randomly, but followed standardized protocols. Benefitting from the participation of 89 citizen scientists, we collected and analyzed approximately 18,000 specimens from over 280 individual fly traps. An in-depth comparison with a biodiversity survey conducted more than 30 years ago revealed significant changes in species abundance and community composition, underscoring the potential of *Drosophila* as a versatile model for monitoring biodiversity loss. Moreover, since *Drosophila* communities consist of species ranging from wild and local to highly synanthropic with global distributions, they provide a powerful system for assessing the degree of anthropogenic impact on ecosystems. Taken together, our study demonstrates that long-term monitoring of *Drosophila*, empowered by citizen science, represents an effective and scalable approach for tracking biodiversity change in a rapidly transforming world.

## Data availability

The full documentation of the data analysis can be found at https://github.com/capoony/UrbanDrosophilaEcology

## Authors contribution

Following the Contributor Role Taxonomy (CRediT; https://credit.niso.org/) MK: Conceptualization, Methodology, Software, Formal analysis, Investigation, Data Curation, Writing – Original Draft, Visualization, Supervision, Project administration, Funding acquisition; SS: Conceptualization, Methodology, Software, Formal analysis, Investigation, Data Curation, Writing – Original Draft; MR: Methodology, Software, Formal analysis, Investigation, Resources, Data Curation, Writing – Original Draft, Writing – Review & Editing; ML: Methodology, Software, Formal analysis, Investigation, Resources, Data Curation, Writing – Original Draft, Writing – Review & Editing; FS: Investigation, Data Curation, Writing – Review & Editing; RQC: Investigation, Data Curation, Writing – Review & Editing; LT: Validation, Investigation, Writing – Review & Editing; NS: Methodology, Validation, Investigation, Writing – Review & Editing; EH: Conceptualization, Methodology, Formal analysis, Investigation, Resources, Writing – Original Draft, Writing – Review & Editing, Supervision

## Acknowledgements

We are deeply indebted and sincerely grateful to the 89 highly dedicated and motivated citizen scientists who contributed the sampling material presented and analyzed in this study. We also thank the Horizon Europe-funded FAIRiCUBE project and consortium (grant agreement No 101059238) for financially supporting this project, and in particular the coordinators, Stefan Jetschny and Katharina Schleidt, for their dedicated guidance. Our gratitude extends to Heimo Rainer, PI of the FAIRiCUBE project at the Natural History Museum Vienna (NHMW), and to Susanna Ioni for their continuous support. We further thank our colleagues at the Central Research Laboratories for assistance with consumables, and the porters at the NHMW for their invaluable help in handling traps – receiving filled ones from citizen scientists and distributing new empty ones. Without the generous contributions of all these individuals, and many more at the NHMW, this project would not have been possible.

## Supporting Information

### Supporting Tables

https://docs.google.com/spreadsheets/d/1p7wvKQCp-Gny8_07CcZTKqa4cR5reTNAxvbv8JZBe7w/edit

### Supporting Figures

**Figure S1.**
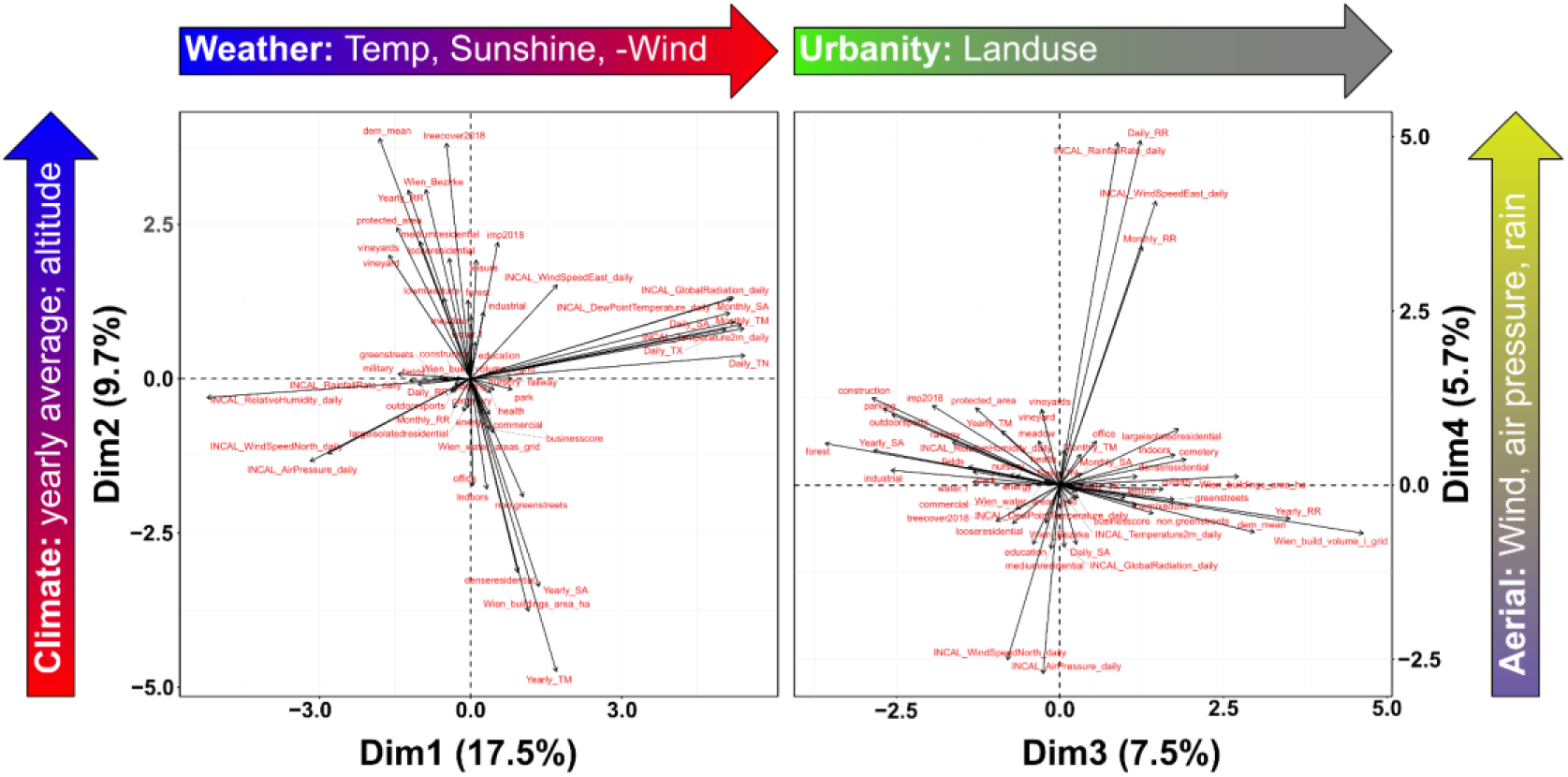
Principal Component Analysis (PCA) of environmental factors. Biplots display the loadings of individual environmental variables along the first two principal components (Dim.1 vs. Dim.2, left) and the third and fourth components (Dim.3 vs. Dim.4, right). Arrows beside each axis summarize the most influential environmental factors contributing to the corresponding dimensions, providing a visual overview of key gradients represented by each principal component.

**Figure S2.**
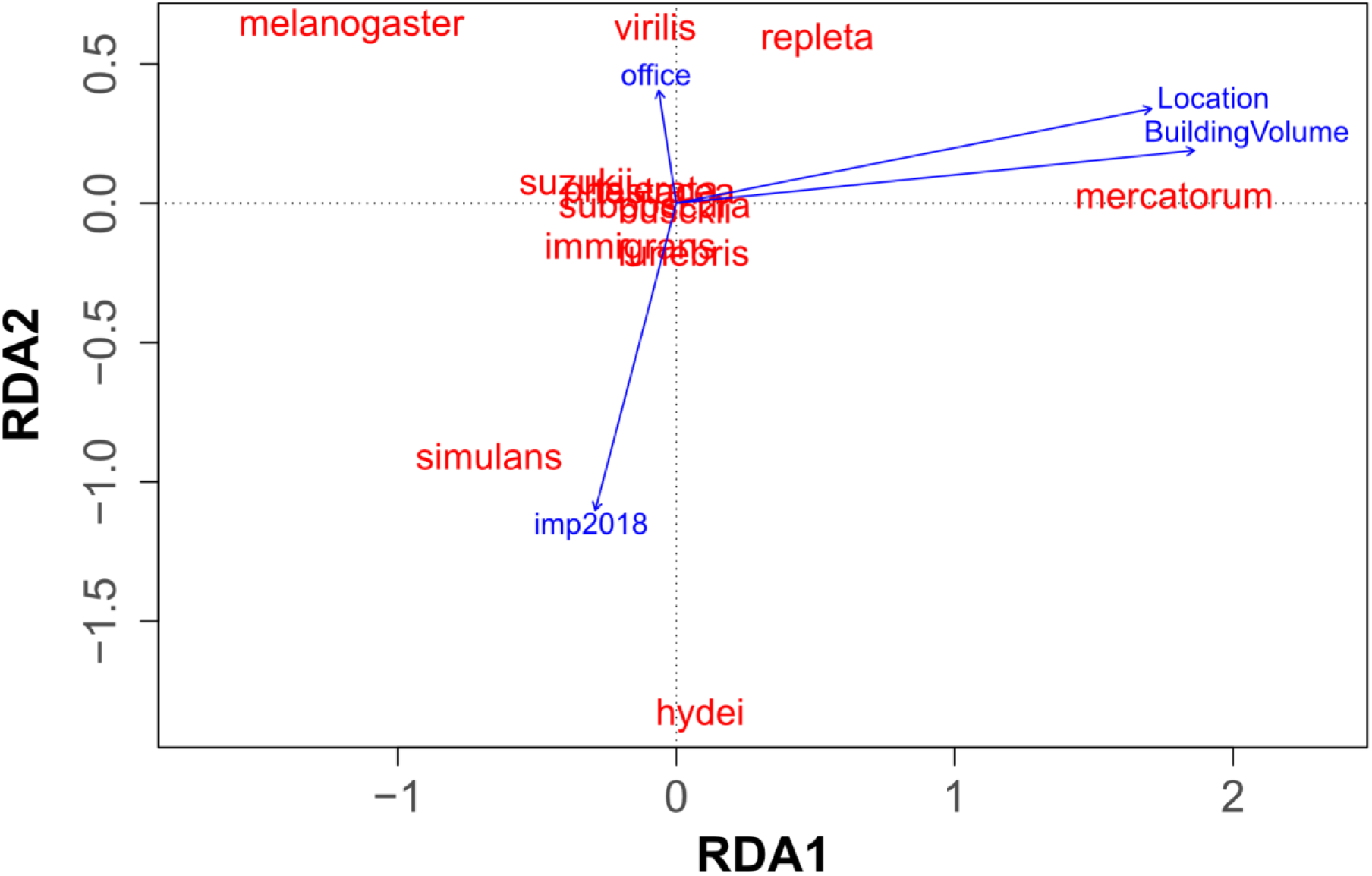
Redundancy Analysis (RDA) of collapsed abundance data. RDA plot showing the first two canonical axes similar to Figure 4B. Red labels indicate the positions of each species in RDA space, and blue arrows represent correlations with four environmental variables (see Table S2) selected by forward regression.

## Notes

### Competing Interest Statement

The authors have declared no competing interest.

