## Supporting Information – Figures for "Ecology and temporal dynamics of urban *Drosophila* species communities as potential indicators of biodiversity decline"

### Supporting Tables

<https://docs.google.com/spreadsheets/d/1p7wvKQCp-Gny8_07CcZTKqa4cR5reTNAxvbv8JZBe7w/edit>

### Supporting Figures

**Figure S1. Principal Component Analysis (PCA) of environmental factors.** Biplots display the loadings of individual environmental variables along the first two principal components (Dim.1 vs. Dim.2, left) and the third and fourth components (Dim.3 vs. Dim.4, right). Arrows beside each axis summarize the most influential environmental factors contributing to the corresponding dimensions, providing a visual overview of key gradients represented by each principal component.

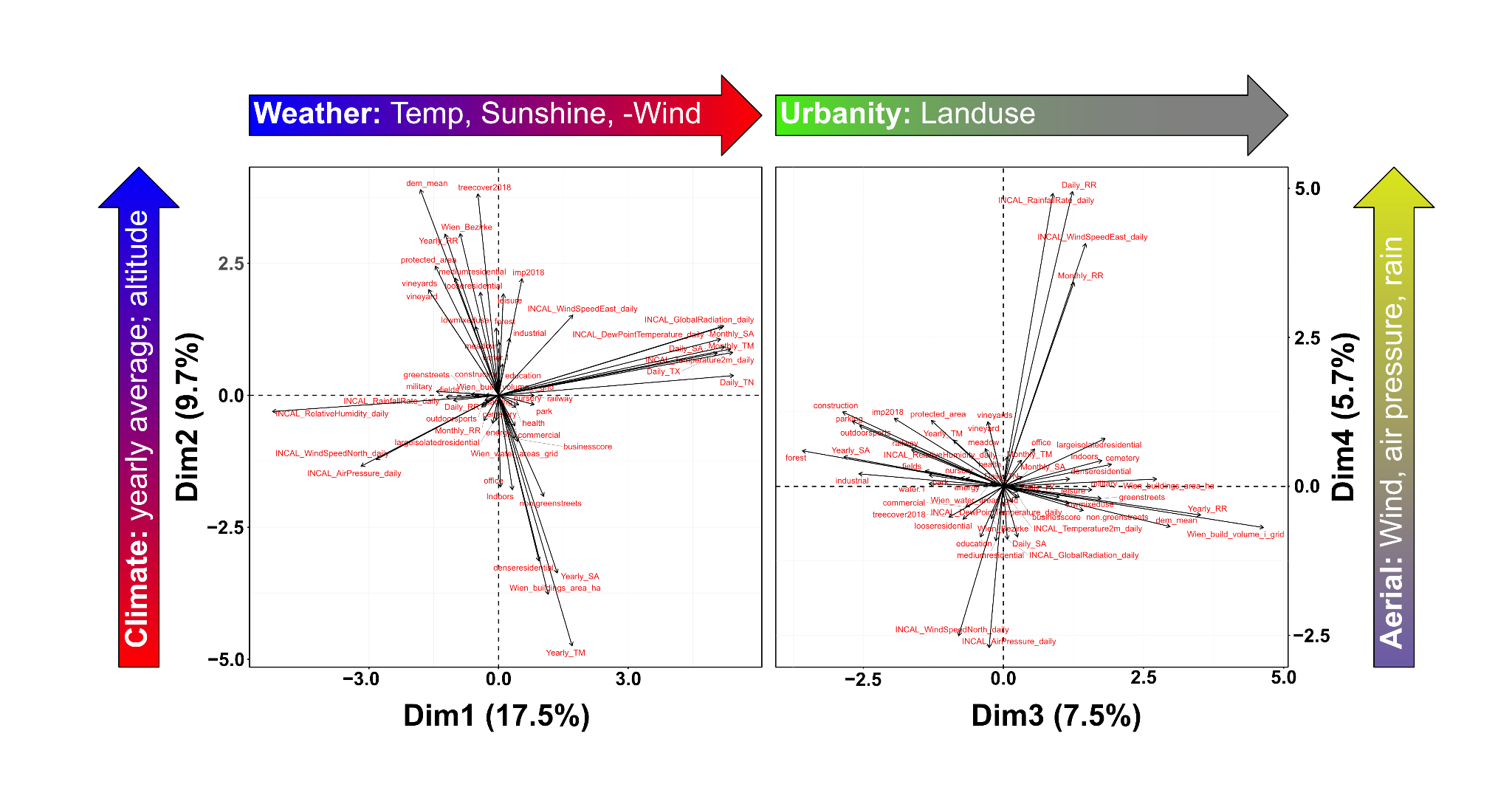

**Figure S2.** **Redundancy Analysis (RDA) of collapsed abundance data**. RDA plot showing the first two canonical axes similar to Figure 4B. Red labels indicate the positions of each species in RDA space, and blue arrows represent correlations with four environmental variables (see Table S2) selected by forward regression.

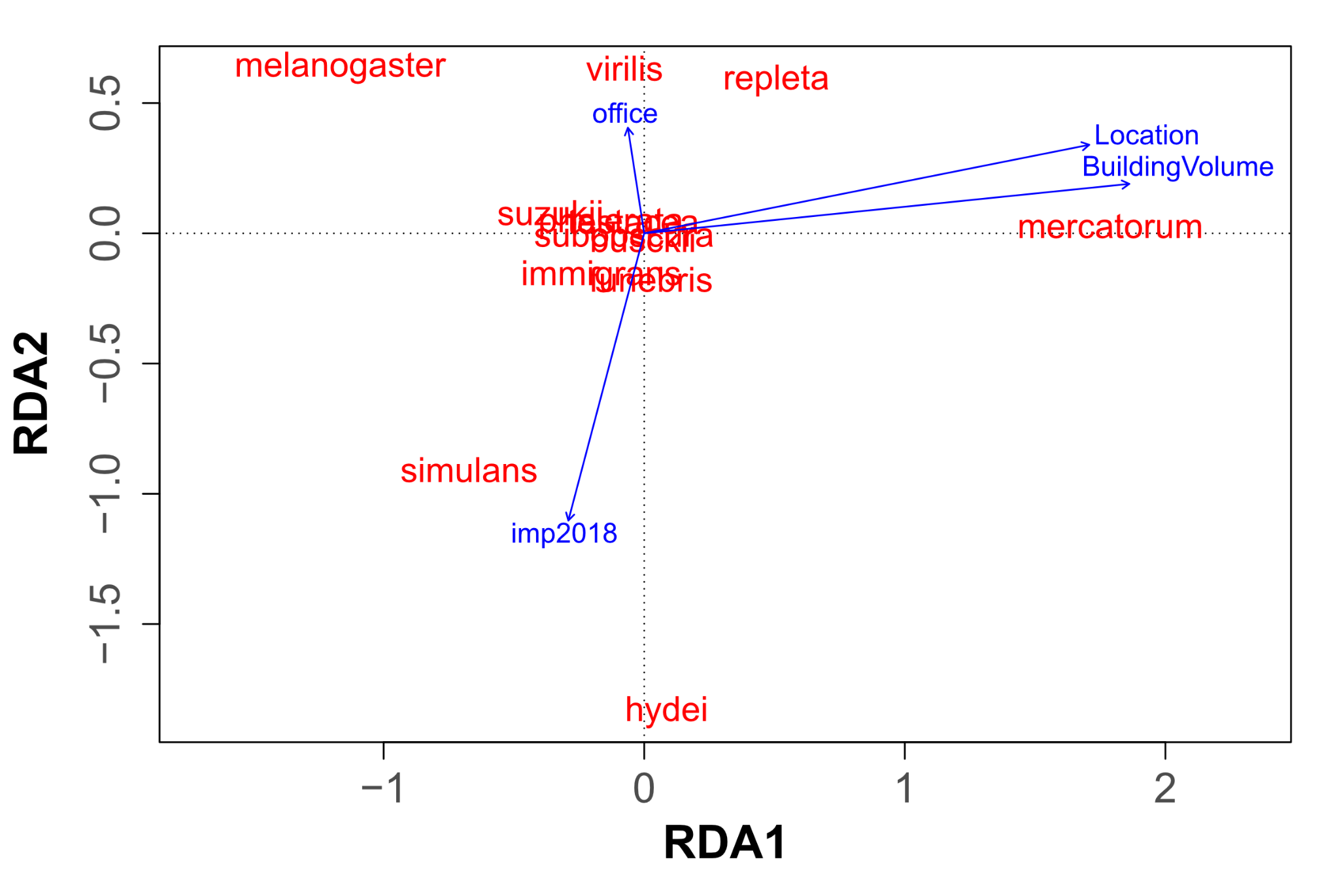
